# Differential transcriptomic responses to heat stress in surface and subterranean diving beetles

**DOI:** 10.1101/2021.12.12.470823

**Authors:** Perry G. Beasley-Hall, Terry Bertozzi, Tessa M. Bradford, Charles S. P. Foster, Karl Jones, Simon M. Tierney, William F. Humphreys, Andrew D. Austin, Steven J. B. Cooper

## Abstract

Subterranean habitats are generally very stable environments, and as such evolutionary transitions of organisms from surface to subterranean lifestyles may cause considerable shifts in physiology, particularly with respect to thermal tolerance. In this study we compared responses to heat shock at the molecular level in a geographically widespread, surface-dwelling water beetle to a congeneric subterranean species restricted to a single aquifer (Dytiscidae: Hydroporinae). The obligate subterranean beetle *Paroster macrosturtensis* is known to have a lower thermal tolerance compared to surface lineages (CT_*max*_ 38°C *cf*. 42-46 °C), but the genetic basis of this physiological difference has not been characterized. We experimentally manipulated the thermal environment of 24 individuals to demonstrate that both species can mount a heat shock response at high temperatures (35°C), as determined by comparative transcriptomics. However, genes involved in these responses differ between species and a far greater number were differentially expressed in the surface taxon, suggesting it can mount a more robust heat shock response; these data may underpin its higher thermal tolerance compared to subterranean relatives. In contrast, the subterranean species examined not only differentially expressed fewer genes in response to increasing temperatures, but also in the presence of the experimental setup employed here alone. Our results suggest *P. macrosturtensis* may be comparatively poorly equipped to respond to both thermally induced stress and environmental disturbances more broadly. The molecular findings presented here have conservation implications for *P. macrosturtensis* and contribute to a growing narrative concerning weakened thermal tolerances in obligate subterranean organisms at the molecular level.

## 1 INTRODUCTION

The transition to an obligate subterranean lifestyle can cause massive shifts in an organism’s biology (1), from the acquisition of classic troglomorphies such as the elongation of appendages for sensing in an aphotic environment (2), to changes in lesser-studied traits including circadian rhythm (3,4), reproductive biology (5), respiration (6), the number of larval instars (7,8) and chemosensation (9,10). These changes have been attributed to the stark difference between subterranean and surface (hereafter epigean) habitats. While epigean environments can vary immensely over both time and space, subterranean environments, such as cave systems, generally possess high environmental stability with respect to light levels, temperature, humidity, and nutrient availability (11–13). Animals adapted to these environments might therefore be particularly well suited for the assessment of responses to future climate change scenarios, particularly with respect to their thermal tolerances and responses to increasing temperatures. Indeed, such habitats have been labelled as undervalued natural laboratories for biological studies of global change (14).

A near-universal response to temperature-induced stress across the tree of life–and therefore a method by which thermal tolerance can be gauged–is the heat shock response (hereafter HSR), which involves the synthesis of heat-shock proteins (hereafter HSPs). HSPs include those proteins that are expressed constitutively under non-stressful conditions, called heat shock cognates, or those only induced when organisms are exposed to thermal extremes, during which they assist in stabilising and refolding proteins at risk of denaturation. An inducible HSR has been observed in almost all organisms studied to date, and the proteins involved in this response, as well as the response itself, are highly conserved among different domains of life (15). An estimated 50 to 200 genes are involved in the HSR, the most significantly induced of which are HSPs (16). However, there are exceptions to this rule: a lack of an inducible HSR has been documented in a wide range of species, largely those that occupy very stable thermal environments such as Antarctic marine habitats (17,18).

Knowledge of the HSR in organisms from thermally stable *subterranean* habitats, including their associated inducible HSPs and at which temperatures this response might be activated, is scarce, with only a few studies devoted to invertebrate taxa (14,19–25). However, invertebrates overwhelmingly contribute to the biodiversity of subterranean habitats compared to vertebrates (26,27). The bulk of existing studies on the thermal tolerance of subterranean invertebrates suggest such taxa can withstand temperatures above those they would encounter in nature and that they have not lost the HSR. Only a small number of these studies have examined the heat shock response directly (28) and tend to focus on thermal tolerances gauged through survival experiments. Moreover, a better understanding of responses to current climate change predictions for subterranean animals has been identified as a fundamental question in subterranean biology given emerging conservation issues associated with their respective ecosystems (29). To address this knowledge gap, here we make use of genomic data from Australian representatives of a group of aquatic invertebrates containing both epigean and subterranean lineages.

The Yilgarn Craton in central Western Australia (WA) houses a diverse subterranean diving beetle fauna belonging to two tribes, Bidessini and Hydroporini (Dytiscidae). While epigean species can be found practically continent-wide, subterranean taxa are isolated in calcrete aquifers (hereafter calcretes) associated with ancient palaeodrainage systems in the region. These calcretes are completely devoid of light and animals contained within them are assumed to have little to no access to air above the water’s surface (6). Each calcrete houses between one and three dytiscid species, but at the time of writing only around a quarter of the ~200 known calcretes have been sampled (30–32). Nonetheless, lineages in both tribes are known to have made independent yet parallel, repeated transitions into underground habitats from epigean ancestors during the late Miocene to early Pleistocene, likely in response to continental aridification (32–35). In each case these transitions have involved the loss of eyes, pigment, and wings (30) as well as the gain of a remarkable ability to respire directly from water (6). Preliminary evidence suggests that subterranean members of these lineages are less tolerant of thermal extremes compared to epigean relatives (36), but the molecular mechanisms underlying such tolerances–and genomic changes associated with a subterranean transition in these animals more broadly–remain unknown. An investigation into the potential link between the above-mentioned mode of respiration, oxygen delivery, and heat tolerance is also lacking (37).

In the present study, we focus on two members of the Hydroporini: *Paroster nigroadumbratus* (Clark), an epigean species endemic to South Australia, and the subterranean *Paroster macrosturtensis* (Watts & Humphreys) found exclusively in a single calcrete at Sturt Meadows in the Yilgarn region of WA. As subterranean dytiscids likely descended from only a handful of epigean lineages, meaningful comparisons can be made between these taxa despite their distributions being geographically disjunct and their divergence *ca*. 15 Mya (32). A recent study showed that *P. macrosturtensis* has a reduced upper critical thermal maximum (CT_max_) of 38.3°C compared to other epigean dytiscids (42–44.5°C) (36), mirroring previous results for other cave beetle species (23). These findings suggest *P. macrosturtensis* is unlikely to reach its thermal critical maximum under current climate change predictions. However, given the thermal stability in its environment, it remains unknown as to whether *P. macrosturtensis* might have a modified HSR compared to its epigean relatives and if exposure to high temperatures may nonetheless induce significant stress in this subterranean species. Here, we present transcriptomic data from individuals of *P. macrosturtensis* and *P. nigroadumbratus* subjected to varying degrees of heat shock following the results of Jones *et al*. (36). In the present study we specifically aimed 1) to generate a high-quality, near-complete reference transcriptome for *P. nigroadumbratus* and 2) using this dataset, characterise and compare the HSR of *P. nigroadumbratus* and *P. macrosturtensis*, specifically with respect to which genes are differentially expressed and the conditions under which this occurs.

## 2 MATERIALS AND METHODS

### 2.1 Taxon sampling, experimental design, and cDNA sequencing

Beetle specimens (*P. macrosturtensis*, n=11; *P. nigroadumbratus*, n=13) were sourced as described in a previous study and subjected to heat stress using an aquarium setup described therein (36).Specimens were placed in one of three groups: in a controlled-temperature cabinet at 25°C (hereafter control), in vials within the experimental setup at 25°C, and in vials within the experimental setup ramped to 35°C. The control temperature of 25°C was selected by (36) to reflect the approximate average groundwater temperatures of the aquifer that *P. macrosturtensis* is found in. Following treatment, the individuals were placed in liquid nitrogen for RNA sequencing immediately after exposure to control temperatures and thermal extremes. RNA extractions were performed using single whole bodies prior to the synthesis and sequencing of barcoded cDNA samples. Quality control of sequence data was performed using Trim Galore with default settings v.0.4.1 (http://bioinformatics.babraham.ac.uk/projects/trim_galore). More information regarding specimen collection and husbandry, experimental design, cDNA sequencing, and phylogenetic analysis can be found in the electronic supplementary material.

### 2.2 Assembly of *Paroster nigroadumbratus* reference transcriptome

A reference transcriptome for *P. nigroadumbratus* was *de novo* assembled using Trinity v.2.5.1 using default settings (38) and derived from unpublished raw RNA-seq data from (39). Resulting transcriptome completeness was validated using BUSCO v.5 (40,41), and functional annotation was performed using TransDecoder v.5.5.0 (42) and the Trinotate v.3.2.1 pipeline (38), which employs SQLite (http://sqlite.org/index.html), BLAST v.2.7.1 (43), and HMMER v.3.2 (http://hmmer.org). Redundancy within the transcriptome assembly was reduced by retaining only the longest isoform for each Trinity gene identifier, following (10), ensuring differential expression analysis was performed at the Trinity ‘gene’ level rather than that of isoforms. This subsetted dataset was used in downstream analyses, though we note this approach limited our ability to examine alternative splice variants.

### 2.3 Differential expression and Gene Ontology enrichment analysis

The above reference transcriptome was indexed and raw reads generated in this study from both *P. macrosturtensis* and *P. nigroadumbratus* were quasi-mapped to it and normalised using Salmon v.1.1.0 (44). Differential expression analysis was performed using edgeR v.3.32.1 (45) alongside Gene Ontology (GO) enrichment analysis, executed using Trinity helper scripts. Differential expression and enrichment/depletion of GO terms was gauged using two comparisons per species (Figure 1). First, for each species the group from the experimental setup at 25°C was compared to the control group outside of the setup at 25°C (hereafter comparison 1). Second, for each species the group in the experimental setup at 35°C was compared to the group in the experimental setup at 25°C (hereafter comparison 2). A diagrammatic representation of these comparisons is shown in Figure 1. These comparisons were designed to gauge the impact of the presence of the experimental setup alone and an increase in temperature, respectively. A full description of this process can be found in the electronic supplementary material.

**FIGURE 1.**
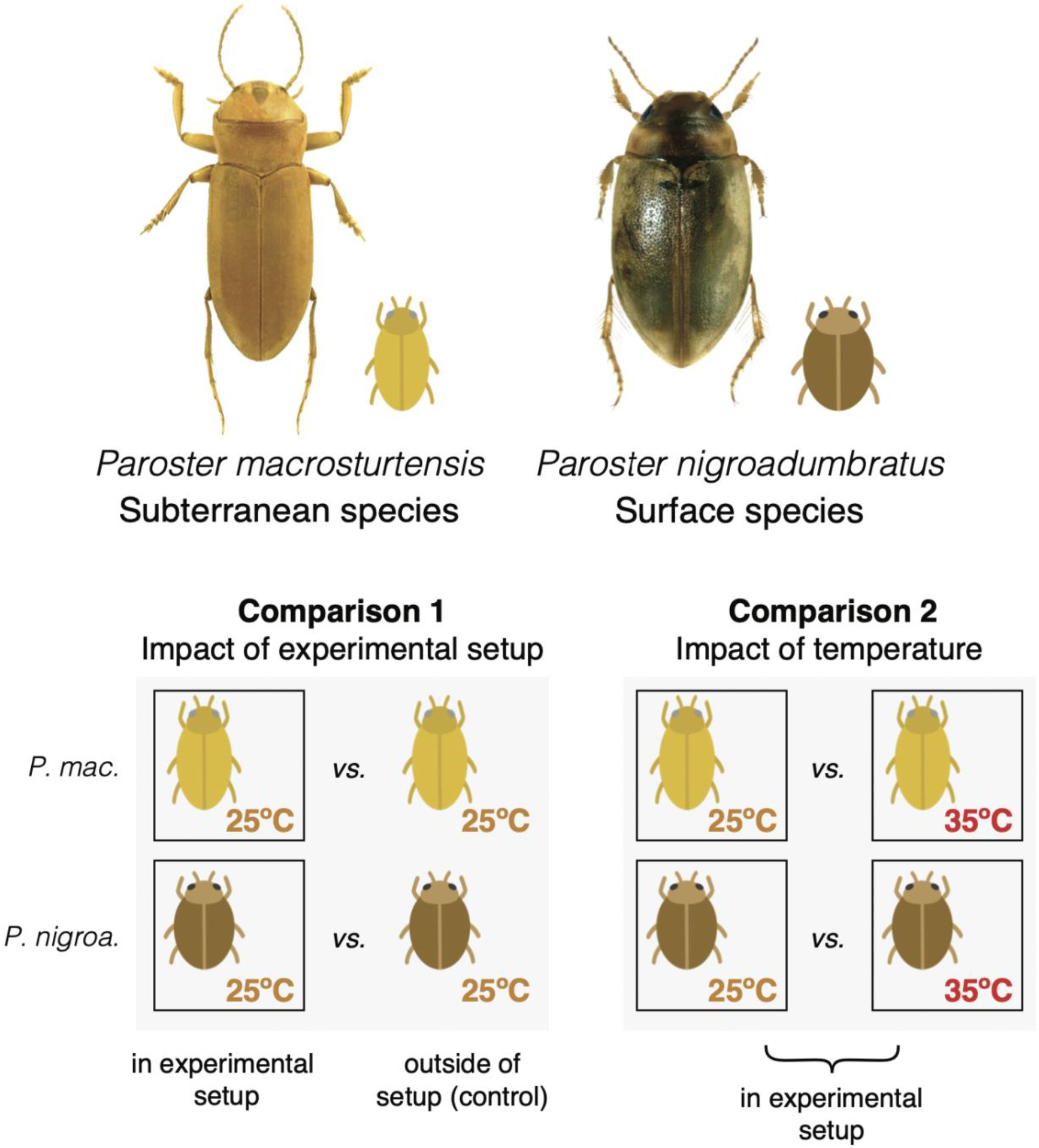
*Paroster* beetles included in this study and the experimental design used to assess differential gene expression associated with thermal extremes alone, as opposed to solely the presence of the experimental setup, following (36). Only intraspecific comparisons were made when assessing differential gene expression in our analyses; interspecific comparisons were made post-hoc. Photographs by Chris Watts and Howard Hamon.

### 2.4 Phylogenetic analysis of putative heat shock proteins

Putative heat shock protein-encoding genes were aligned with coleopteran orthologs sourced from the OrthoDB v.10.1 catalogue (46) and GenBank to confirm their identity (Table S1). Phylogenetic inference for each HSP family was performed in RAxML v.8.2.12 (47).

## 3 RESULTS

### 3.1 A high-quality reference transcriptome for *Paroster nigroadumbratus* enables the characterisation of genes involved in the heat shock response

Here we present a high-quality, near-complete transcriptome for the epigean beetle *P. nigroadumbratus*. This dataset consisted of 75,045,266 paired-end reads (72,266,264 following quality control measures), 84% of which were incorporated into 134,246 *de novo* assembled transcripts representing 60,683 unique Trinity ‘genes’and 41,979 predicted ORFs. According to the assessment using BUSCO, this transcriptome was 87.66% complete with respect to complete core arthropod genes and 96.74% complete when considering partial genes. Of these transcripts, 47,810 were able to be functionally annotated using the Trinotate pipeline. Subsetting our predicted peptide dataset to include only the longest isoforms per Trinity gene identifier, allowing us a proxy with which to perform our downstream analyses at the gene level, resulted in 14,897 predicted ORFs (with 11,609, or ~77%, having some level of annotation).

### 3.2 Differential expression analysis reveals distinct expression profiles associated with the heat shock response in *Paroster*

We compared expression profiles between members of the same species, subjected to different conditions, to assess genes differentially expressed in response to the presence of the experimental setup employed here (comparison 1) or an increase in temperature in that setup (comparison 2) (Figure 2). Parallels and contrasts between *P. nigroadumbratus* and *macrosturtensis* were then assessed post-hoc. Differential expression analysis using edgeR identified a total of 723 differentially expressed (DE) genes in the epigean *P. nigroadumbratus* samples and 157 in the subterranean *P. macrosturtensis*, with the two species exhibiting complex and markedly different expression profiles (Figures 2, 3, S2, S3). *P. nigroadumbratus* consistently differentially expressed a greater number of genes than *P. macrosturtensis*: 147 and 67 genes were differentially expressed in comparison 1 and 89 and 51 genes were differentially expressed in comparison 2, respectively. The presence of the experimental setup and an increase in water temperature caused both species to differentially express genes involved in the heat shock response. Contrasting expression profiles with respect to the genes involved were not only observed between the two species (as above) but also in the response of each species to these two different stressors (Figure 2).

**FIGURE 2.**
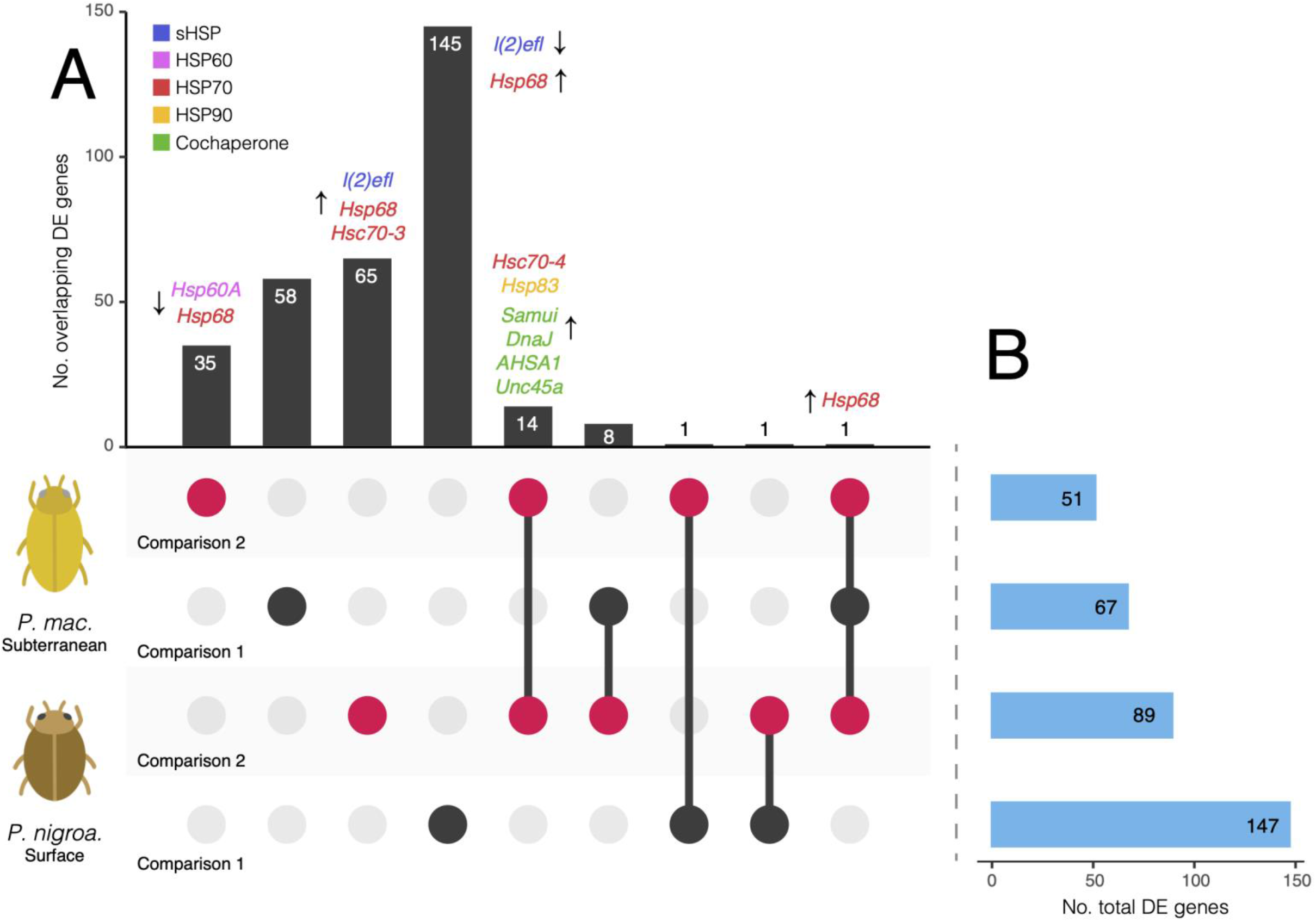
Differentially expressed (DE) genes in *Paroster macrosturtensis* and *nigroadumbratus* in comparison 1 (the experimental setup used in this study relative to the control, black circles) or comparison 2 (35°C within the setup relative to 25°C in the setup, red circles). Comparisons are ordered by total number of DE genes. A) DE genes shared (circles linked by lines) or unique to (unlinked circles) each comparison per species, summed in the bar graph above. Up- or downregulated HSPs or HSP cochaperones are shown for each group. The gene *Hsp68* being named more than once in different groups refers to separate transcripts sharing the same putative annotation (see Table S2). B) Total DE genes for each species under different conditions. HSP gene names were sourced from Trinotate annotations and orthology was validated using phylogenetic analysis (Figure 4).

**FIGURE 3.**
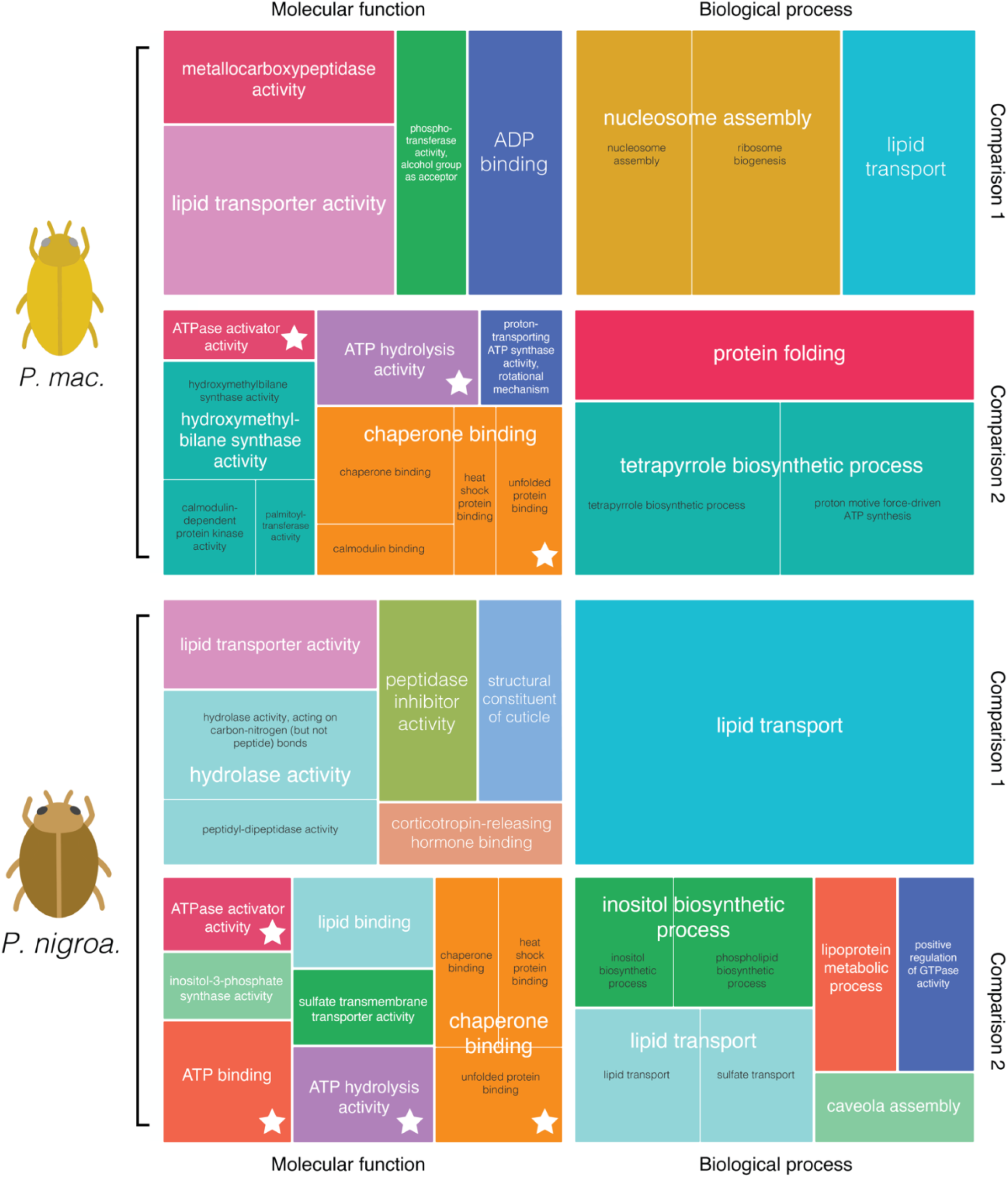
REVIGO (48) treemaps showing enriched Gene Ontology (GO) terms associated with the differential expression of genes shown in Figure 2. Treemaps are labelled as comparison 1 (the experimental setup used in this study relative to the control) or 2 (35°C within the setup relative to 25°C in the setup). The size of squares is proportional to the *p*-value associated with differential expression of respective genes. Similar GO terms share a colour and are represented in white text by the largest square per group. GO terms associated with genes involved in the heat shock response are indicated by a star.

In comparison 1, the surface species *P. macrosturtensis* significantly upregulated a HSP gene putatively annotated as *Hsp68*, encoding a major heat shock protein in the HSP70 family, relative to the control. *Paroster nigroadumbratus* upregulated the HSP *Hsp68* and downregulated the sHSP *l(2)efl* in the experimental setup relative to the control. None of the HSP transcripts differentially expressed by *P. nigroadumbratus* in response to the presence of the experimental setup only were differentially expressed by *P. macrosturtensis* or by *P. nigroadumbratus* in comparison 2.

In comparison 2, both species upregulated the HSP genes *Hsc70-4* and *Hsp83* as well as the cochaperones *Samui, DnaJ, AHSA1*, and *Unc45a* relative to groups at the lower temperature of 25°C. Both *P. nigroadumbratus* and *macrosturtensis* also upregulated separate Trinity “genes” annotated as *Hsp68* each at 35 °C relative to the 25 °C treatment (Fig. 2), likely representing closely related loci that are yet to be comprehensively characterised in the beetles. Additional differentially expressed genes unique to each species at 35°C relative to the 25°C treatment included: 1) downregulation of the HSPs *Hsp60A* and *Hsp68* in *Paroster macrosturtensis*, and 2) the upregulation of the sHSP *l(2)efl*, the HSP *Hsp68*, and the HSP cognate *Hsc70-3* in *P. nigroadumbratus*. Other annotated genes potentially involved in the HSR, such as *Hsc70-2, Hsc70-5, Trap1*, and *Hsp90b1*, were not differentially expressed in either species (Figure 4). A full list of differentially expressed genes shared between (or unique to) the two beetle species under different conditions is available in Table S2. Proteins unrelated to the HSR, yet widely differentially expressed in our dataset (i.e., with reoccurring annotations across different Trinity ‘genes’) included those involved in the transport of lipids and nutrient storage, such as vitellogenin and apolipophorins.

**FIGURE 4.**
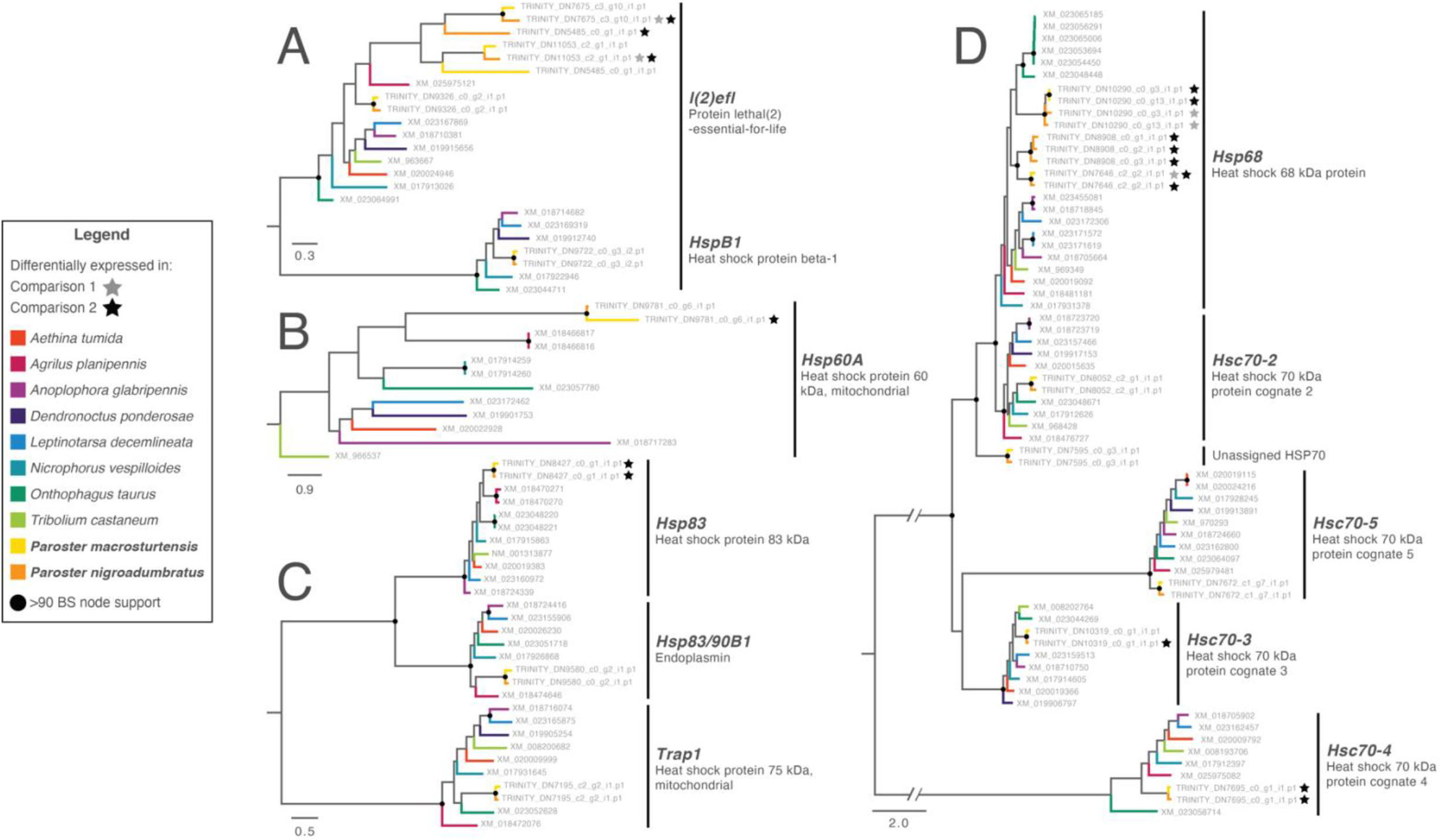
Phylogenies of heat shock Trinity ‘genes’ inferred using RAxML to validate the identity of putative HSP orthologs in *Paroster* species. Trees are as follows: A) sHSP family, B) HSP60, C) HSP90, D) HSP70. Scale bar is in substitutions/site; *BS* = bootstrap node support. Tip labels show transcript names/GenBank accession numbers. Tip names for *P. macrosturtensis* are shared with the reference *P. nigroadumbratus* transcript reads were assembled against. Tip names with stars specifically refer to genes differentially expressed in the presence of the experimental setup alone relative to the control (comparison 1) or at 35°C relative to 25°C in the experimental setup (comparison 2).

Gene Ontology enrichment analysis enables a high-order approximation of the functional consequences of differentially expressed genes. We did not observe any consistent depletion of GO terms associated with downregulated genes, but enriched terms were reflective of our differential expression results above (Figure 3). In both species, terms associated with the HSR were enriched in association with an increased temperature at 35°C (e.g., ATPase activator activity [GO:0001671], chaperone binding [GO:0051087], and unfolded protein binding [GO:0051082]). The most common remaining Gene Ontology terms included lipid transporter activity (GO:0005319), nutrient reservoir activity (GO:0045735), metal ion binding (GO:0046872), and zinc ion binding (GO:0008270).

### 3.3 Phylogenetic analyses of HSPs

Our HSP nucleotide alignments consisted of 12 sequences across 1,743 bp (HSP60 family), 67 sequences across 3,669 bp (HSP70 family), 29 sequences across 3,741 bp (HSP90 family) and 23 sequences across 867 bp (sHSP family). Sequences clustered by gene with strong node support in our phylogeny (Figure 4). All sequences from the *Paroster* species examined here were recovered as nested within these clades, confirming the orthology of annotated HSPs, with the exception of a transcript present in both species and inferred as sister to the *Hsp68+Hsc70-2* clades. This gene was not differentially expressed in response to heat shock in our dataset.

## 4 DISCUSSION

Here, we have comprehensively characterised the heat shock response at the molecular level in the subterranean diving beetle *Paroster macrosturtensis* and one of its surface-dwelling relatives. Using a near-complete reference transcriptome for *P. nigroadumbratus*–the first such dataset for a member of the Hydroporinae–we performed differential expression and GO enrichment analysis to explore genes putatively involved in the HSR. Our results demonstrate that both the epigean *P. nigroadumbratus* and subterranean *P. macrosturtensis* have an inducible HSR, in agreement with implications of previous survival experiments for the genus (36). Putative orthologs of HSP cochaperones, sHSP, HSP60, HSP70, and HSP90 genes were accounted for in our differential expression analysis. However, the conditions under which this response is activated differs between species, and *P. macrosturtensis* notably differentially expressed just over half of the number of genes compared to *P. nigroadumbratus* in response to a rise in temperature.

### 4.1 Heat shock-induced gene expression in *Paroster*

HSPs identified as differentially expressed in this study support past results for beetle species and other insects more broadly. A major trend in our results concerned the upregulation of HSP70 genes at high temperatures, particularly *Hsp68;* HSP70s are highly expressed in response to heat shock in other beetle species, including cave-adapted subterranean taxa (28,49,50), and work in concert with sHSPs and HSP90s (51–54). In addition to the heat shock proteins, we also observed the upregulation of the heat shock cognates *Hsc70-3* and *Hsc70-4* and putative cochaperones *Tsc2, Samui, DnaJ, AHSA1, Unc45a* at 35°C relative to the 25°C treatment. To our knowledge there has been no documentation of a coleopteran heat shock cognate being upregulated in response to increasing temperature, though evidence exists for the parasitic wasp *Pteromalus*, in which *hsc70* is induced by heavy metal poisoning and starvation in addition to thermal extremes (55). Heat shock cognates are also upregulated during diapause in silkworm eggs (56) and young bumble bee queens (57), potentially playing a cryoprotective role in these species. Cochaperones are less well characterised in insects, but evidence for their upregulation in response to heat shock has been documented in hemipterans and ants (58–60). We observed the downregulation of several heat shock proteins in both species in the presence of the experimental setup relative to the control, and at high temperatures relative to the 25°C treatment. Both sHSPs and HSP70s have been documented as being downregulated during periods of heat stress in other insects, e.g. in silk moths (61). In *P. nigroadumbratus* this was restricted to the sHSP *l(2)efl* and cochaperone *Tsc2* in the experimental setup-only comparison. In contrast, the HSPs *Hsp60A* and *Hsp68* were downregulated *in P. macrosturtensis* at high temperatures relative the 25°C treatment.

### 4.2 Expression profiles reflect differing thermal tolerances

Our molecular data mirrors previously documented reduced thermal tolerances in subterranean insects such as *P. macrosturtensis*. The species differentially expressed far fewer genes in response to 35°C relative to the 25°C treatment compared to *P. nigroadumbratus;* similarly reduced numbers of differentially expressed genes have also been associated with lower thermal tolerances in other organisms such as fish (62), lizards (63), rotifers (64), red algae (65), and plants (66,67), though we note the inverse (or alternatively, no clear pattern) has been observed in a number of cases, potentially reflecting lower levels of stress as opposed to an inability to mount a HSR (68,69).

In keeping with the above findings, *P. macrosturtensis* also differentially expressed far fewer genes than its epigean counterpart in response to the presence of the experimental setup alone relative to the control (Figure 2). HSPs are known to be involved in responding to a wide range of stressors (70–72), and the involvement of such genes is not surprising in stress unrelated to temperature; individuals being moved into the experimental setup employed here may have induced stress from handling, for example. The greater number of genes differentially expressed by *P. nigroadumbratus* in this scenario may suggest *P. macrosturtensis* is potentially less able to robustly respond to ambient stressors more broadly (i.e., environmental disturbances). Such a scenario is supported by past work showing subterranean species are sensitive to ambient stressors under otherwise non-stressful temperatures (73), and being an epigean species, *P. nigroadumbratus* is presumably exposed to far more dramatic environmental fluctuations (in addition to more variable temperatures) on a regular basis than a subterranean species such as *P. macrosturtensis*. We also note that in the presence of the experimental setup relative to the control, *P. macrosturtensis* upregulated the same *Hsp68*-annotated Trinity ‘gene’ implicated in responses to heat-induced stress in both species (Figure 2), whereas HSPs differentially expressed by *P. nigroadumbratus* under the same conditions did not overlap with those in other groups.

### 4.3 Heat shock and the climatic variability hypothesis

The dataset we present here adds to a growing body of knowledge concerning the HSR in organisms that inhabit thermally stable environments. Central to discourse on this topic is the climatic variability hypothesis, which posits that the thermal tolerance of a taxon is positively correlated with its temperature ranges encountered in nature (74). This hypothesis implies species from extremely stable thermal environments can no longer tolerate temperature extremes, and has been demonstrated in a wide variety of organisms that have either lost or possess a reduced HSR, such as cnidarians (75), limpets (76), amphipods and sea stars (77), and midges (78). In contrast, species that inhabit areas with a broader range of climatic conditions would be expected to be more robust in the face of environmental fluctuations (79). While *P. macrosturtensis* does have a lower thermal tolerance compared to *P. nigroadumbratus*, in line with the above hypothesis, it nonetheless has retained a HSR at high temperatures per our transcriptomic data. Similar studies have shown certain groundwater-dwelling organisms display an inducible HSR in response to conditions far warmer than they would encounter in nature (20,21). The HSR of these species, as well as *P. macrosturtensis*, might be retained at such high temperatures for a variety of reasons, including the fact that the species has not occupied its respective environments for a sufficient length of time in evolutionary terms for their HSR to be lost, e.g. via adaptive processes or a relaxation of purifying selection (36). The latter scenario is plausible as *P. macrosturtensis* is also known to have retained the ability to detect light despite inhabiting an aphotic environment for over ~3 million years (80).

### 4.4 Conservation implications

While the retention of a HSR in both species examined here supports the physiological findings of Jones *et al*. (36), almost half (4 out of 10 assayed) of the *P. macrosturtensis* cohort did not survive 24 hours after heat shock in that study. It therefore remains to be seen if the species can tolerate such extremes in the long term. Indeed, even cave beetles considered stenothermal–those that are only capable of surviving within an extremely narrow temperature range–have retained the HSR (23), but nonetheless cannot survive at extreme temperatures for long periods (>7 days) compared to epigean relatives (25,28). Threatening processes that *P. nigroadumbratus* and *macrosturtensis* are both at risk of experiencing in their fragile habitats might impact the latter species far more negatively as a result.

Temperature rises of up to 5°C by the end of the century compared to pre-industrial levels may occur in central Western Australia per current climate change projections (81). Water temperatures in aquifers are generally cooler and more stable than, but are nonetheless coupled with, conditions above-ground, and are also predicted to warm as regional temperatures increase (82–84). The subterranean habitat of these insects is therefore unlikely to shield them from the impacts of a warming world. The fact that *P. macrosturtensis* appears to be unable to mount as robust a HSR compared to *P. nigroadumbratus*, and therefore may experience a significantly higher amount stress compared to epigean species in the face of high temperatures, has conservation implications for the understudied fauna of the Australian Yilgarn and beyond.

Datasets such as these are especially pertinent for subterranean invertebrates found in the Yilgarn region—including *P. macrosturtensis* and its subterranean relatives, in addition to crustaceans such as isopods and amphipods—as the groundwater in their calcrete habitats is heavily utilised for water extraction by industry (85,86). As short-range endemics to the extreme, such species are not only at risk of habitat degradation via climate change, but from the direct intersection of shallow aquifers with e.g. mining activities and through the drawdown of groundwater beneath calcretes at greater depths (87). In addition to reflecting the reduced thermal tolerances of Australian subterranean dytiscids, the molecular data we presented here for *P. macrosturtensis* also suggests a potentially weaker response in the face of other environmental disturbances unrelated to temperature. These factors have the potential to render *P. macrosturtensis* more vulnerable to both of the above threatening processes compared to epigean relatives, with implications for subterranean fauna more broadly. An increased knowledge of the assumed fragility of Australian subterranean invertebrates in the face of these stressors is therefore crucial for informing future conservation management plans for these animals and their fragile habitats.

## 5 CONCLUSIONS

Our findings demonstrate the reduced thermal tolerance of the subterranean species *P. macrosturtensis* compared to its epigean relatives is reflected, and further clarified by, transcriptomic data. While our data are supported by past physiological evidence that demonstrated *P. macrosturtensis* could survive at high temperatures, albeit not to the limits of epigean species (36), the present study adds a new layer to this narrative. *P. macrosturtensis* might possess increased mortality in the face of high temperatures because the species differentially expresses far fewer genes in response to heat shock compared to the epigean relative *P. nigroadumbratus*, suggesting it may be unable to mount as robust a heat shock response. While *P. macrosturtensis* might be able to survive at temperatures far above those it encounters in nature for short periods, as detailed by Jones *et al*. (36), our results suggest the species also experiences a weaker transcriptomic response to factors unrelated to temperature (i.e. the presence of the experimental setup employed here) relative to *P. nigroadumbratus*. As such, *P. macrosturtensis* may not be as well-equipped to survive higher temperatures and other threatening processes, such as disturbances to surrounding groundwater, in the long term compared to surface-dwelling members of *Paroster*. Future work in this system will ideally assess a far greater number of dytiscid species to further explore the trends we observe here. As the present study did not consider the role of isoforms in the heat shock response of these animals, broader studies could examine these responses to heat stress at a finer scale by conducting differential expression analysis on the transcript, as opposed to gene, level.

## ACKNOWLEDGEMENTS

We thank Flora, Peter, and Paul Axford for providing access to the Sturt Meadows calcrete and accommodation at the Sturt Meadows pastoral property. We thank Chris Watts (South Australian Museum) for helping with identification and collecting of surface dytiscids, providing beetle images, and for laying the foundation for the subterranean beetle research with WFH and Remko Leijs. We thank Rae Humphreys for assistance with field collections of *P. macrosturtensis* and for being a wonderful host to stygofauna catchers over many years. Finally, we would like to thank two anonymous reviewers for helping to improve the quality of the manuscript.

## AUTHOR CONTRIBUTIONS AND COMPETING INTERESTS STATEMENT

PGBH, KJ, and SJBC conceived the study. KJ performed experiments exposing beetles to heat stress. PGBH analysed the data and wrote the manuscript. TB and CSPF provided guidance regarding bioinformatic analyses. ST performed molecular laboratory work. TMB, KJ, WFH, and ADA helped to draft the manuscript. All authors edited the manuscript and approved the final draft. Funding for this project was provided by an Australian Research Council Discovery grant (DP180103851) to SJBC, WFH, ADA and TB. The authors declare that they have no competing interests.

## DATA AVAILABILITY STATEMENT

All raw RNA-seq data for *Paroster macrosturtensis* and *P. nigroadumbratus* used in differential expression analyses in this study are available via NCBI under BioProject PRJNA783065 (individual Sequence Read Archive accessions for samples SRR17023302-SRR17023315). HSP transcripts assembled from raw *P. macrosturtensis* RNA-seq data and used in phylogenetic analysis are available via GenBank (accession numbers summarized in Table S1). The reference transcriptome of *P. nigroadumbratus* (including all isoforms) and the annotated, subsetted dataset (only including longest isoform per Trinity ‘gene’) are available via FigShare (DOI: 10.25909/17169191).

