## Supplementary Material for "Differential transcriptomic responses to heat stress in surface and subterranean diving beetles"

**Supplementary Materials and Methods**

***Taxon sampling and experimental design***

Adult beetle specimens were sourced as described in previous study (1) that subjected specimens to heat stress. A subset of individuals was assayed in survival experiments, described in (1). The remainder were euthanized immediately after exposure to heat stress for RNA sequencing, which we describe in the present study. We briefly reiterate the methodology used in (1) below.

*Paroster macrosturtensis* samples (3.6–4.1 mm long, n = 11) were collected from a 3.5km^2^ region of the Sturt Meadows calcrete in WA via existing mineral exploration boreholes between five and 11 m in depth. Epigean *P. nigroadumbratus* samples (3.2mm long, n = 13) were collected from ephemeral pools in the Mount Crawford region of South Australia. The sex of specimens was unable to be determined without immobilising individuals, which had the potential to bias levels of stress experienced by the beetles. Individuals were kept in constant temperature cabinets at between 24 and 26ºC either in a 12:12h light-dark cycle (epigean species) or in complete darkness (subterranean species) for at least seven days prior to subsequent experimentation to minimise the effects of variable thermal history and stress during capture and transport. Stygobiotic species were fed twice a week on pieces of fresh blackworm (*Lumbricus*), while epigean species were fed live blackworms *ad libitum*. Beetles were either retained in this constant temperature environment and euthanized in liquid N_2_ (control treatment) or moved to an experimental setup within a temperature-controlled water bath containing pure reverse osmosis (RO) water (epigean species) or RO water with a salinity level mimicking calcrete microhabitats (subterranean species). The experimental setup consisted of plastic vials containing one individual beetle each. Vials with subterranean beetles had a layer of mesh and fabric around the circumference to allow water circulation, which could otherwise limit access to oxygen (and therefore thermal tolerance) due to the aquatic, cutaneous nature of respiration in *P. macrosturtensis* (2). Beetles were kept at 25ºC in this environment for 4.5 hours to determine if they could survive the experimental setup (25ºC treatment) or allowed to acclimate to 25ºC for 1 hour before the temperature was increased by 0.05ºC every minute until it reached 35ºC (35ºC treatment). Ramping rates were informed by field measurements regarding maximum rates of water temperature increases (1). In all cases individuals were euthanized in liquid N_2_ immediately after this time period.

***cDNA synthesis and sequencing***

RNA extractions of control and heat-treated individuals were performed using single whole bodies per treatment. Extractions were performed using an AllPrep DNA/RNA Micro kit (Qiagen, Chadstone, VIC), initially homogenizing in lysis buffer containing β-mercaptoethanol plus a 5 mm steel bead on a Mixer Mill (Retsch, Haan, Germany). Lysates were processed following the manufacturer’s instructions and RNA was eluted from the RNeasy MinElute spin column in 25 µl RNAse-free water. Concentration of the RNA extract was determined using a Quantus Flourometer (Promega, Madison, Wisconsin, US) and RNA integrity was analysed on a TapeStation (Agilent Technologies, Santa Clara, California, US). Library preparation was performed using a Nugen Universal Plus mRNA-Seq Library Preparation Kit (Tecan, Männedorf Switzerland), following manufacturer’s instructions. All the barcoded cDNA samples were pooled and 100 bp paired-end sequencing was outsourced to the Australian Genome Research Facility (AGRF) on an Illumina NovaSeq 6000 S4 lane. The reference transcriptome for *P. nigroadumbratus* presented here was derived from previously obtained raw RNA-seq data in (3).

***Quality control of sequence data***

Trim Galore v.0.4.1 ([bioinformatics.babraham.ac.uk/projects/trim_galore](https://www.bioinformatics.babraham.ac.uk/projects/trim_galore)), a wrapper that implements Cutadapt (4) and FastQC ([bioinformatics.babraham.ac.uk/projects/fastqc](http://bioinformatics.babraham.ac.uk/projects/fastqc)), was used for quality control of our raw reads and the removal of residual sequencing adapters, with default settings for paired Illumina reads. This protocol was applied to both the raw reads from control/heat-treated samples and those intended for reference transcriptome assembly.

**P. nigroadumbratus *reference transcriptome assembly and annotation***

Following quality control and adapter removal, Trinity v.2.5.1 (5) was used for *de novo* transcriptome assembly of our raw reads using default settings. Bowtie v.1.1.2 (6) was used to examine RNA-Seq read representation in the assembly by aligning raw reads to the assembled transcriptome. Reference transcriptome completeness was validated using the gVolante implementation of BUSCO v.5 (7,8) with default settings, searching against the Arthropoda database.

Functional annotation of the transcriptome was performed using TransDecoder v.5.5.0 (9) to predict coding regions in transcripts, Trinotate v.3.2.1 (5) and SQLite (<http://sqlite.org/index.html>) for database integration, and BLAST v.2.7.1 (10) and HMMER v.3.2 (<http://hmmer.org>) for homology searches and the identification of protein domains. BLASTX and BLASTP searches were conducted against the Swiss-Prot database v.2021_02 (11) for Trinity transcripts and TransDecoder-predicted proteins, respectively. These searches were performed with an e-value of 1-e6 to capture more distantly matching proteins and *-max_hsps* and *-max_target_seqs* both set to 1 to limit our results to the best hit per query by e-value. While the use of the latter parameter has been subject to scrutiny as top hits are selected based on sequence order in the database in the case of equivalent matches, this behaviour appears to occur in datasets with an extremely high proportion of gaps (12,13), which does not apply to our study. The Pfam database v.34.0 (14) was used in HMMER searches to identify protein domains with default settings. Finally, we subsetted the ORFs inferred by TransDecoder to only include the longest isoform per Trinity gene. This was performed such that each ‘gene’ was only represented once in the dataset, e.g. there would be a single occurrence of gene identifier “TRINITY_DN9248_c1_g5”, represented by its longest isoform.

***Differential expression and Gene Ontology enrichment analysis***

Salmon v.1.1.0 (16) was used to index our annotated *P. nigroadumbratus* reference transcriptome and quasi-map our raw reads from control/heat-treated samples against it with an automatic inference of library type and 100 bootstrap replicates. Reads were normalised with the trimmed mean of M-values normalization method (TMM). TMM read counts for each replicate were visualised using principal components analysis consisting of a total of 13 *P. nigroadumbratus* replicates and 11 for *P. macrosturtensis* in our downstream differential expression analysis (Figure S1).

The Trinity helper scripts for edgeR v.3.32.1 (17) were used for differential expression analyses and enrichment of Gene Ontology (hereafter GO) terms based on annotations from the *P. nigroadumbratus* reference transcriptome. Abundance estimates from Salmon were first transformed for use in edgeR prior to running differential expression analysis with default settings (*abundance_estimates_to_matrix.pl, run_DE_analysis.pl*). Differential expression analysis results were then analysed with a *p*-value cut-off for FDR of 0.05 and a fold-change of 1. This analysis included the enrichment/depletion of GO terms by supplying Trinity gene lengths and GO terms for each annotation (18) and compared 1) beetles within the experimental setup at 25ºC to beetles outside of the setup at the same temperature, and 2) beetles at 25º to those at 35ºC, both within the experimental setup.

Significantly up- and downregulated genes shared between different treatments were identified using a custom Venn diagram webtool ([bioinformatics.psb.ugent.be/webtools/Venn](http://bioinformatics.psb.ugent.be/webtools/Venn/)) and visualised with UpSetR v.1.4.0 (19). Enrichment of GO terms was visualised with REVIGO (20), which summarises terms by clustering them and retrieving a meaningful subset using similarity measures. Enriched GO terms were supplied alongside FDR values associated with enrichment/depletion and the REVIGO analysis was performed using an allowed similarity of 0.9. Resulting tree maps were produced with values set to abs(log10(*p*-value)) to ensure plots reflected the significance of enrichment of each term.

***Phylogenetic analysis of putative heat shock proteins***

Genes annotated as putatively encoding heat shock proteins consisted of members of three of the four insect families of HSP, including small heat shock proteins (sHSP), HSP60, HSP70, and HSP90. Transcripts were extracted based on annotations containing the case-insensitive terms *heat shock*, *HSC*, *HSP*, or *chaperone*. To ensure the inclusion of as many genes as possible for which annotation might have been incomplete (e.g. certain HSP chaperones), we also included transcripts putatively involved in the HSR that were differentially expressed in our dataset yet not captured in the above search. The resulting candidate set of transcripts was checked against the UniProt database to confirm the function of the proteins they encoded, and the identities of transcripts were then validated using BLASTN searches restricted to the Coleoptera against the NCBI nr database. These transcripts were used as a reference to filter raw reads from each *P. macrosturtensis* 35ºC replicate using BWA v.0.7.17 (21) and SAMtools v.1.9 (22). Filtered *P. macrosturtensis* reads (excluding singletons) were then pooled and mapped against reference transcripts using the Read Mapper tool in Geneious Prime (v.2021.2.2, [geneious.com](http://geneious.com)) with Low Sensitivity presets. Consensus sequences for *P. macrosturtensis* were produced from these alignments with a 0% (majority) threshold.

Coleopteran orthologs of the above genes were retrieved from OrthoDB v.10.1 (23) with LOC identifiers of the top BLASTN hit as queries. mRNA transcripts of protein sequences in question were obtained from GenBank, representing sequences from *Aethina tumida* Murray (Polyphaga: Cucujoidea), *Agrilus planipennis* Fairmaire (Polyphaga: Buprestoidea), *Anoplophora glabripennis* (Motschulsky) (Polyphaga: Chrysomeloidea), *Dendroctonus ponderosae* (Hopkins) (Phytophaga: Curculionoidea), *Leptinotarsa decemlineata* Say (Polyphaga: Chrysomeloidea), *Nicrophorus vespilloides* Herbst (Polyphaga: Staphylinoidea), *Onthophagus taurus* (Schreber) (Polyphaga: Scarabaeoidea), and *Tribolium castaneum* (Herbst) (Polyphaga: Tenebrionoidea) alongside our two species of interest (summarised in Table S1). Alignments were computed using MUSCLE v.3.8.425 (24), checked by eye to ensure all transcripts were in frame, and trimmed to exclude portions of GenBank sequences outside of open reading frames. Partitions were inferred using PartitionFinder v.2.1.1 (25), with a resulting partitioning scheme by codon position. Phylogenetic analysis of HSPs was performed using maximum likelihood with RAxML v.8.2.12 (26) via the CIPRES Science Gateway (27) with default settings and 1000 bootstrap replicates.

**Supplementary Figures and Tables**

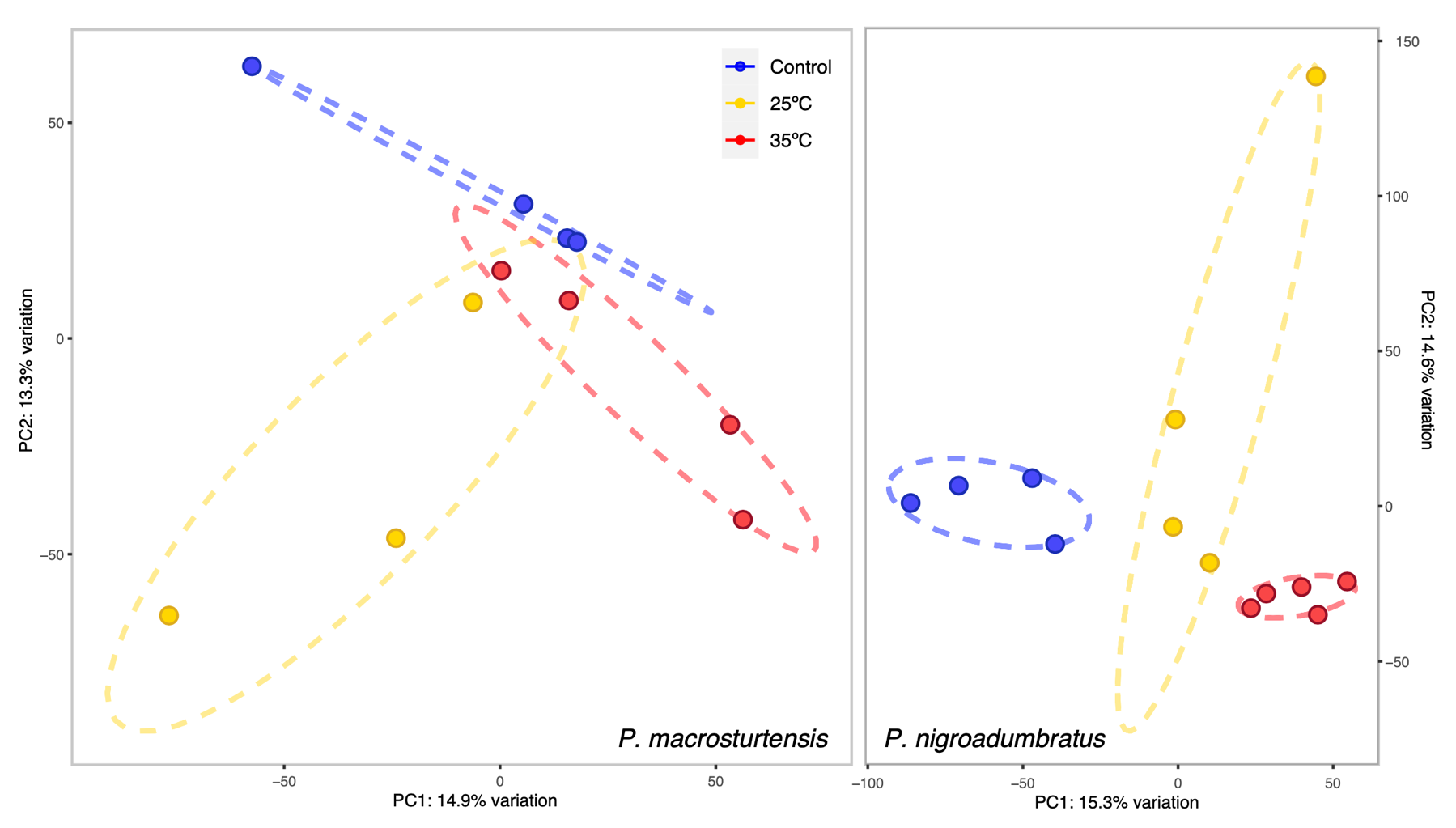

**FIGURE S1** Principal components analysis of TMM-normalized read counts for *P. nigroadumbratus* and *P. macrosturtensis* individuals subjected to different temperature treatments following (1). Abbreviations: 25ºC treatment outside of experimental setup (control), 25ºC treatment over 4.5 hours within experimental setup (25ºC), ramping treatment from 25ºC to 35ºC over ~4 hours (35ºC).

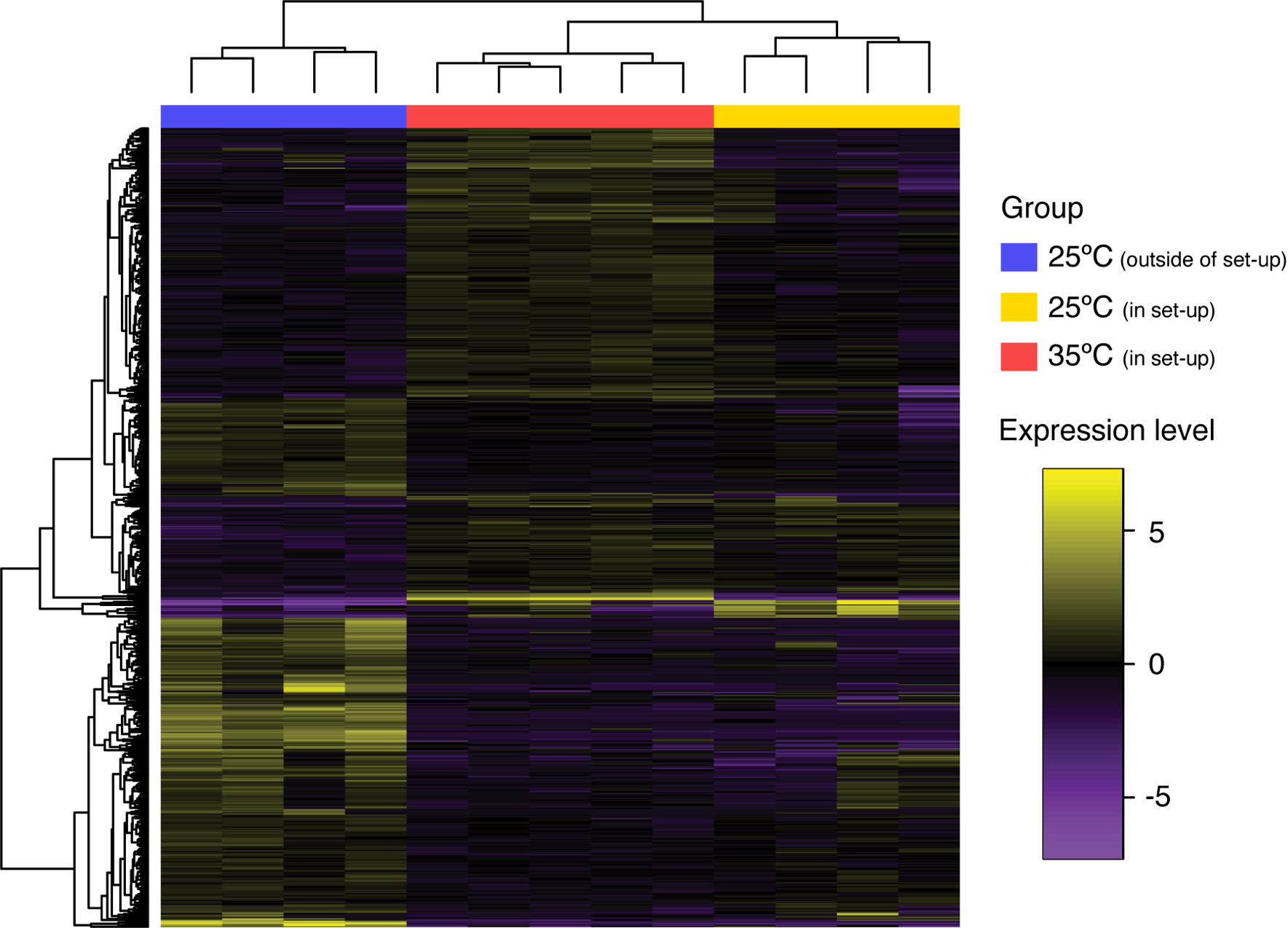

**FIGURE S2** Heatmap of the 723 differentially expressed genes in *P. nigroadumbratus* individuals at different temperature treatments from edgeR with a *p*-value cut-off for FDR of 0.05 and a fold-change of 1.

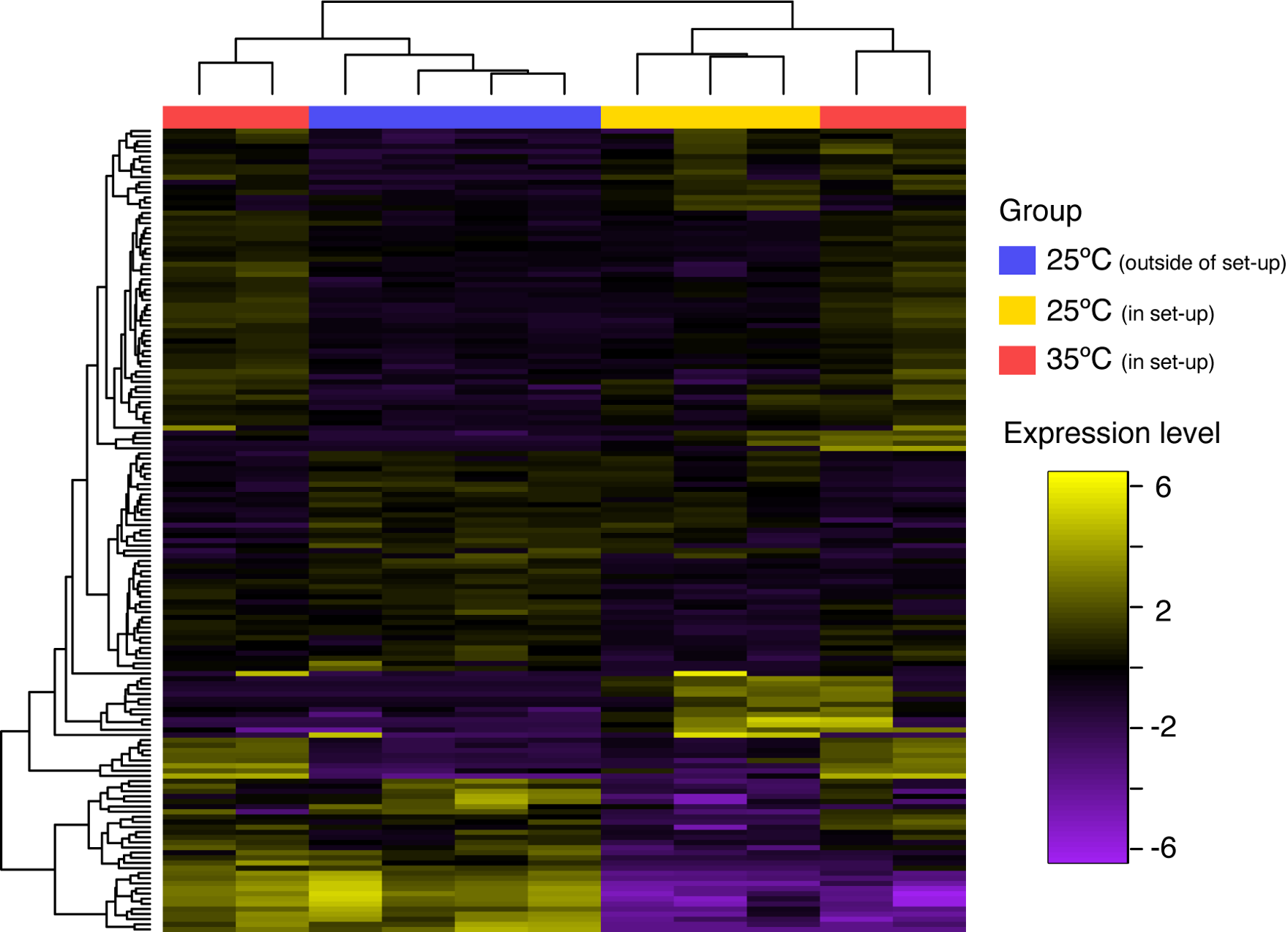

**FIGURE S3** Heatmap of the 157 differentially expressed genes in *P. macrosturtensis* individuals at different temperature treatments from edgeR with a *p*-value cut-off for FDR of 0.05 and a fold-change of 1.

**TABLE S1** NCBI accessions and BioProject numbers corresponding to other members of the Coleoptera included in the present study for phylogenetic analysis. As raw *P. macrosturtensis* data were mapped against the reference transcriptome presented here and do not originate from an assembled, annotated transcriptome, we also provide the NCBI accessions for the single HSP genes of that species used for the phylogeny in Figure 4.

| **Gene name** | **Species** | **GenBank accession number(s)** | **BioProject/reference** |
| --- | --- | --- | --- |
| Unidentified HSP70 mRNA | *Paroster macrosturtensis* | OL803878 | - |
| *Hsc70-2* | *Aethina tumida* | XM_020015635 | PRJNA361278 |
|  | *Agrilus planipennis* | XM_018476727 | PRJNA343475 |
|  | *Anoplophora glabripennis* | XM_018723719, XM_018723720 | PRJNA167479, PRJNA348318 |
|  | *Dendroctonus ponderosae* | XM_019917153 | PRJNA360270 |
|  | *Leptinotarsa decemlineata* | XM_023157466 | PRJNA420356 |
|  | *Nicrophorus vespilloides* | XM_017912626 | PRJNA339573 |
|  | *Onthophagus taurus* | XM_023048671 | PRJNA419349 |
|  | *Paroster macrosturtensis* | OL803874 | - |
|  | *Tribolium castaneum* | XM_968428 | PRJNA15718 |
| *Hsc70-3* | *Aethina tumida* | XM_020019366 | PRJNA361278 |
|  | *Anoplophora glabripennis* | XM_018710750 | PRJNA348318 |
|  | *Dendroctonus ponderosae* | XM_019906797 | PRJNA360270 |
|  | *Leptinotarsa decemlineata* | XM_023159513 | PRJNA420356 |
|  | *Nicrophorus vespilloides* | XM_019714605 | PRJNA339573 |
|  | *Onthophagus taurus* | XM_023044269 | PRJNA419349 |
|  | *Paroster macrosturtensis* | OL803875 | - |
|  | *Tribolium castaneum* | XM_008202764 | PRJNA15718 |
| *Hsc70-4* | *Aethina tumida* | XM_020009792 | PRJNA361278 |
|  | *Agrilus planipennis* | XM_025975082 | PRJNA343475 |
|  | *Anoplophora glabripennis* | XM_018705902 | PRJNA348318 |
|  | *Leptinotarsa decemlineata* | XM_023162457 | PRJNA420356 |
|  | *Nicrophorus vespilloides* | XM_017912397 | PRJNA339573 |
|  | *Onthophagus taurus* | XM_023058714 | PRJNA419349 |
|  | *Paroster macrosturtensis* | OL803876 | - |
|  | *Tribolium castaneum* | XM_008193706 | PRJNA15718 |
| *Hsc70-5* | *Aethina tumida* | XM_020019115, XM_020024216 | PRJNA361278 |
|  | *Agrilus planipennis* | XM_025979481 | PRJNA343475 |
|  | *Anoplophora glabripennis* | XM_018724660 | PRJNA348318 |
|  | *Dendroctonus ponderosae* | XM_019913891 | PRJNA360270 |
|  | *Leptinotarsa decemlineata* | XM_023162800 | PRJNA420356 |
|  | *Nicrophorus vespilloides* | XM_017928245 | PRJNA339573 |
|  | *Onthophagus taurus* | XM_023064097 | PRJNA419349 |
|  | *Paroster macrosturtensis* | OL803877 | - |
|  | *Tribolium castaneum* | XM_970293 | PRJNA15718 |
| *Hsp60A* | *Aethina tumida* | XM_020022928 | PRJNA866502 |
|  | *Agrilus planipennis* | XM_018466816, XM_018466817 | PRJNA343475 |
|  | *Anoplophora glabripennis* | XM_018717283 | PRJNA348318 |
|  | *Dendroctonus ponderosae* | XM_019901753 | PRJNA846874 |
|  | *Leptinotarsa decemlineata* | XM_023172462 | PRJNA420356 |
|  | *Nicrophorus vespilloides* | XM_017914259, XM_017914260 | PRJNA339573 |
|  | *Onthophagus taurus* | XM_023057780 | PRJNA419349 |
|  | *Paroster macrosturtensis* | OP296523 | - |
|  | *Tribolium castaneum* | XM_966537 | PRJNA15718 |
| *Hsp68* | *Aethina tumida* | XM_020019092 | PRJNA361278 |
|  | *Anoplophora glabripennis* | XM_018705664, XM_018718845, XM_023455081 | PRJNA348318 |
|  | *Leptinotarsa decemlineata* | XM_023171572, XM_023171619, XM_023172306 | PRJNA420356 |
|  | *Nicrophorus vespilloides* | XM_017931378 | PRJNA339573 |
|  | *Onthophagus taurus* | XM_023048448, XM_023053694, XM_023054450, XM_023056291, XM_023065006, XM_023065185 | PRJNA419349 |
|  | *Paroster macrosturtensis* | OL803879, OL803880, OL803881 | - |
|  | *Tribolium castaneum* | XM_969349 | PRJNA15718 |
| *Hsp83* | *Aethina tumida* | XM_020019383 | PRJNA361278 |
|  | *Agrilus planipennis* | XM_018470270, XM_018470271 | PRJNA343475 |
|  | *Anoplophora glabripennis* | XM_018724339 | PRJNA348318 |
|  | *Leptinotarsa decemlineata* | XM_023160972 | PRJNA420356 |
|  | *Nicrophorus vespilloides* | XM_017915863 | PRJNA339573 |
|  | *Paroster macrosturtensis* | OL803883 | - |
|  | *Onthophagus taurus* | XM_023048220, XM_023048221 | PRJNA419349 |
|  | *Tribolium castaneum* | NM_001313877 | Peuß *et al.* (2015) |
| *Trap1* | *Aethina tumida* | XM_020009999 | PRJNA361278 |
|  | *Agrilus planipennis* | XM_018472076 | PRJNA343475 |
|  | *Anoplophora glabripennis* | XM_018716074 | PRJNA348318 |
|  | *Dendroctonus ponderosae* | XM_019905254 | PRJNA360270 |
|  | *Leptinotarsa decemlineata* | XM_023165875 | PRJNA420356 |
|  | *Nicrophorus vespilloides* | XM_017931645 | PRJNA339573 |
|  | *Paroster macrosturtensis* | OL803882 | - |
|  | *Onthophagus taurus* | XM_023052628 | PRJNA419349 |
|  | *Tribolium castaneum* | XM_008200682 | PRJNA15718 |
| *Hsp90B1* | *Aethina tumida* | XM_020026230 | PRJNA361278 |
|  | *Agrilus planipennis* | XM_018474646 | PRJNA343475 |
|  | *Anoplophora glabripennis* | XM_018724416 | PRJNA348318 |
|  | *Leptinotarsa decemlineata* | XM_023155906 | PRJNA420356 |
|  | *Nicrophorus vespilloides* | XM_017926868 | PRJNA339573 |
|  | *Paroster macrosturtensis* | OL803873 | - |
|  | *Onthophagus taurus* | XM_023051718 | PRJNA419349 |
| *HspB1* | *Anoplophora glabripennis* | XM_018714682 | PRJNA348318 |
|  | *Dendroctonus ponderosae* | XM_019912740 | PRJNA360270 |
|  | *Leptinotarsa decemlineata* | XM_023169319 | PRJNA420356 |
|  | *Nicrophorus vespilloides* | XM_017922946 | PRJNA339573 |
|  | *Paroster macrosturtensis* | OL803884 | - |
|  | *Onthophagus taurus* | XM_023044711 | PRJNA419349 |
| *l(2)efl* | *Aethina tumida* | XM_020024946 | PRJNA361278 |
|  | *Agrilus planipennis* | XM_025975121 | PRJNA343475 |
|  | *Anoplophora glabripennis* | XM_018710381 | PRJNA348318 |
|  | *Dendroctonus ponderosae* | XM_019915656 | PRJNA360270 |
|  | *Leptinotarsa decemlineata* | XM_023167869 | PRJNA420356 |
|  | *Nicrophorus vespilloides* | XM_017913026 | PRJNA339573 |
|  | *Paroster macrosturtensis* | OL803885, OL803886, OL803887, OL803888 | - |
|  | *Onthophagus taurus* | XM_023064991 | PRJNA419349 |
|  | *Tribolium castaneum* | XM_963667 | PRJNA15718 |

**TABLE S2** Differentially expressed genes in *Paroster* shared (or unique to) each species examined and whether each gene is up- or downregulated. These data are available as a supplementary file (Table_S2.xlsx) and were used to construct Figure 2.
